# Tuberculosis in the military: Trends in notification and treatment outcomes within the Zimbabwe Defence Forces, Dependants and Communities 2010-2015

**DOI:** 10.1101/382325

**Authors:** Caroline Muringazuva, Philip Owiti, Jeffrey Edwards, Godfrey N Mutetse, Munyaradzi Dobbie, Jasper Chimedza

## Abstract

**Background:** Historically, tuberculosis (TB) has been responsible for significant disease burden among the military both during peace and conflict. A routine review of Zimbabwe Defence Force (ZDF) data showed 36% of reported deaths could be attributed to TB. We conducted a study to determine the TB trends and outcomes among patients managed in ZDF health facilities between 2010-2015.

**Methods:** Retrospective cohort study of TB patients (military and dependents). Data were extracted from ZDF TB registers and analyzed for trends in notification and outcomes. Independent factors associated with unfavourable TB treatment outcomes were modeled using multivariable regression.

**Results:** Of the total 1298 TB patients, 84% were males, median age 37 years and 92% from Army facilities. Ninety three percentage had pulmonary TB, 87% were new patients and 68% HIV co-infected (97% on antiretroviral therapy[ART]). Number of TB cases reduced two-fold between 2010-2015 (317 vs 115). Treatment outcomes remained relatively stable with overall treatment success of 81%, 9.9% deaths, 0.2% loss to follow up, 2.2% treatment failure, 6.6% not evaluated. Clients who were HIV-positive and not on ART were 3.81 times likely to have unfavourable outcome.

**Conclusion:** This is the first study of TB in an African defence force showed decreasing trends in notified TB cases. Though treatment success was comparable over time, it still fell below international targets. Being HIV-positive (even with ART) was associated with increased unfavourable outcomes. Continued monitoring, evaluation and increased support of the TB programme within this high risk population is recommended.

## Introduction

Tuberculosis (TB) has become the leading infectious cause of death in Africa and a major global health problem(1). In 2016, there were an estimated 10.4 million TB cases globally, 10% of these among people living with HIV (PLWHIV), and 25% in the African region(2). The number of TB deaths is unacceptably high, but with timely diagnosis and treatment, almost all with TB can be cured The disease led to approximately 1.7 million deaths in the same year (1). The global plan to end TB introduced three people-centred targets called the 90-(90)-90 targets: reach 90% of all people who need TB treatment, including 90% of people in key populations, and achieve at least 90% treatment success(2).

The Sustainable Development Goals (SDGs) for 2030 were adopted by the United Nations in 2015 and a key target is to end the global TB epidemic(3). The WHO End TB Strategy, approved by the World Health Assembly in 2014, calls for a 90% reduction in TB deaths and an 80% reduction in the TB incidence rate by 2030(4). Zimbabwe is one of 14 high burden countries for TB, TB/HIV and multidrug-resistant TB (MDR-TB) in the world (5). The focus of the country is to detect all TB cases early, particularly the bacteriologically positive cases, and provide them with effective treatment in a patient centred approach so as to reduce individual short and long term TB morbidity and mortality, stop TB transmission and reduce or eliminate the risk of development of drug resistance (6). In 2015, TB treatment coverage, defined as the number of new and relapse cases that were notified and treated, divided by the estimated number of all incident TB cases that occurred in that year, was estimated at 72% while the treatment success rate (the percentage of all notified TB patients who were successfully treated) was 81%. (2) This performance is below the global benchmark target for effective TB care and prevention of at least 90% treatment coverage and 90% treatment success rate (7). In Zimbabwe, TB is closely associated with HIV infection with an estimated 70% of TB cases being HIV co-infected (5).

Historically, TB has been responsible for significant disease burden among the military, during both times of peace and armed conflict (8). This is likely due to a number of factors including: living and working in confined environments with crowding during training and deployment, personnel may be deployed to highly TB endemic areas, and the physiologically stressful activities during training and operations (8). There are few published studies on TB in the military and none from Africa.

The ZDF is composed of the Zimbabwe National Army (ZNA) and the Air Force of Zimbabwe (AFZ). The two services have health facilities which provide weekly surveillance with monthly and quarterly reports to their headquarters, which then forwards the data for analysis to ZDF Headquarters (HQ) Health Services. The military accommodates most of its members and dependants in military camps, offering all health services free of cost. However, with the introduction of medical aid societies, which soldiers were encouraged to join, some members receive health services in out-of-camp facilities.

In the ZDF, routine review of 2016 end-of-year data showed that 36% of all reported deaths could be attributed to TB. However, trends in TB notification and treatment outcomes have not yet been evaluated. It is against this background that we carried out a study to determine the frequency of TB notification, clinical characteristics and treatment outcomes among all patients seen in the ZDF health facilities from 2010-2015.

## Materials and Methods

### Study Design

We conducted a cohort study utilizing routinely collected programmatic data.

### General setting

Zimbabwe is a lower middle income country in southern Africa. The country is divided into ten administrative provinces and 62 districts. The capital city is Harare and other major cities include Bulawayo, Gweru, Kadoma, Kwekwe, Masvingo and Mutare. The population of Zimbabwe is estimated to be 16 million with 52% being female (9). The Ministry of Health and Child Care provides health coverage in the country and is essentially free of cost.

### The National Tuberculosis Control Programme

The Zimbabwe National TB Control program (NTP) is responsible for control of TB in the country(9). The goals of the NTP are to find at least 90% of new TB cases, provide care for 90% of new TB patients, decrease mortality due to TB, decrease the challenges placed upon families and communities, and to ultimately eliminate TB(6). The NTP program also oversees TB control in the Uniformed Forces. In the ZDF, TB management is done through the environmental health, the laboratory and the curative department. The country’s NTP program provides commodities, training and all monitoring, evaluation and reporting resources for the military TB program.

In terms of TB Management in Zimbabwe the TB Policy points out that (9):

1. Tuberculosis screening and diagnosis using the Xpert MTB/Rif assay and sputum smear microscopy for follow up is provided free of charge.
2. Chemotherapy for all forms of TB, is provided free of charge in the public health sector, and according to international guidelines.
3. TB services are made available at all levels of the health delivery system and integrated into the primary health care system to ensure efficient case finding and case holding.
4. Collaborative TB/HIV activities are carried out at all levels.
5. All patients on treatment for TB receive directly observed treatment either by a health care worker or a community health care worker.

Zimbabwe has established healthcare services countrywide and well developed HIV/TB programs that are supported by collaborative partners. There are more than 100 TB centres nationally and care is provided free of cost at MoHCC facilities(9) The 2016 Global report indicated that 7imbabwe’s TB treatment success rate was 81% and mortality at 7.2% (7).

There are 19 health facilities in the ZDF located in all 10 provinces of Zimbabwe. The military health facilities serve the military members and their dependants. TB diagnosis is being done in 14 of the health facilities.

### Study population

All TB patients registered for TB treatment in the military facilities between 2010 and 2015 were included in the study. Patients included ZDF military personnel and their dependents.

### Data variables, sources of data and data collection

Data was extracted from the ZDF health facility TB registers into a data collection form, entered into EpiData Entry v3.1 (*EpiData Association*, *Odense*, *Denmark*) and cross-checked for consistency. Variables included demographic and clinical characteristics, including type of TB (PTB and EPTB), classification of TB (new or previously treated), HIV status, ART status. Outcomes of treatment were according to standard WHO guidelines (10). Favourable treatment outcome was defined as the aggregate of cure and treatment complete while unfavourable treatment outcome was the aggregate of treatment failure, loss to follow up, died and outcome not recorded (10). Source of data were the TB treatment registers in each of the 14 health facilities in the ZDF.

### Analysis and statistics

Analysis was performed in EpiData Analysis software. Categorical data were analysed and presented in frequencies (and proportion) and comparisons made by the chi-square test. Continuous variables were assessed by medians (interquartile range) and compared by the Mann Whitney U test. Factors associated with unfavourable TB treatment outcomes was assessed at both bivariate and multivariate levels and presented using relative risk and their 95% confidence intervals (CI). Only factors with *P*<0.2 at bivariate level and age and sex were modelled in the log-binomial regression. Level of significance was set at ≤ 5%.

### Ethics approval

Ethics Issues: Permission for the study was granted by the Zimbabwe Defence Forces and National Tuberculosis Control Program Zimbabwe (Zimbabwe NTCP). Ethics approval was obtained from the Medical Research Council of Zimbabwe. As the study involved anonymized secondary data, informed consent was waived.

## Results

**Table 1** shows the demographic and clinical characteristics of tuberculosis patients served by Zimbabwe Defence Forces health facilities, 2010-2015. A total of 1298 TB patients were included into the study, majority (84%) of whom were males and median age 37 years (interquartile range, IQR: 29-48). More patients were from the Army facilities (92%) than the Airforce. The main type of TB was pulmonary TB at 93% while most of the participants were new patients (87%). HIV infection was diagnosed in 68% of the patients, of whom 97% were on antiretroviral therapy.

**Table 1.** Demographic and clinical characteristics of tuberculosis patients served by Zimbabwe Defence Forces health facilities, 2010-2015

| Variable | Category | Frequency | (%) |
| --- | --- | --- | --- |
| Sex | Female | 205 | (15.8) |
|  | Male | 1093 | (84.2) |
| Age (years) | Median [IQR] | 37 | [29-48] |
|  | 0-14 | 2 | (0.2) |
|  | 15-24 | 142 | (11.5) |
|  | 25-34 | 371 | (30.0) |
|  | 35-44 | 289 | (23.4) |
|  | 45-54 | 341 | (27.6) |
|  | 55-64 | 59 | (4.8) |
|  | 65+ | 32 | (2.6) |
| Health Facility | Army facilities | 1198 | (92.3) |
|  | Air force facilities | 100 | (7.7) |
| TB Type | PTB | 817 | (93.3) |
|  | EPTB | 59 | (6.7) |
|  | *Not recorded* | 422 |  |
| TB classification | New | 1119 | (87.3) |
|  | Previously treated | 162 | (12.7) |
|  | *Not recorded* | 17 |  |
| HIV Status | Positive | 829 | (67.6) |
|  | Negative | 398 | (32.4) |
|  | *Not recorded* | 71 |  |
| On ART | Yes | 793 | (96.5) |
|  | No | 29 | (3.5) |
|  | *Not recorded* | 7 |  |
| Province | Harare | 510 | (39.3) |
|  | Bulawayo | 238 | (18.3) |
|  | Midlands | 159 | (12.3) |
|  | Mash West | 173 | (13.3) |
|  | Manicaland | 189 | (14.6) |
|  | Mash Central | 29 | (2.2) |

IQR – interquartile range; PTB – pulmonary tuberculosis; EPTB – extra pulmonary tuberculosis; ART-antiretroviral therapy

Total number of notified TB cases reduced more than two-fold between 2010 and 2015 (**Figure 1**). Treatment outcomes remained relatively unchanged over the six year period (**Figure 2**) with the overall treatment success being 81.1%. Key adverse outcomes were deaths at 9.9%, loss to follow up at 0.2% and not evaluated at 6.6% (**Table 2**). Overall, 19% of patients had unfavourable TB treatment outcomes.

**Fig 1.**
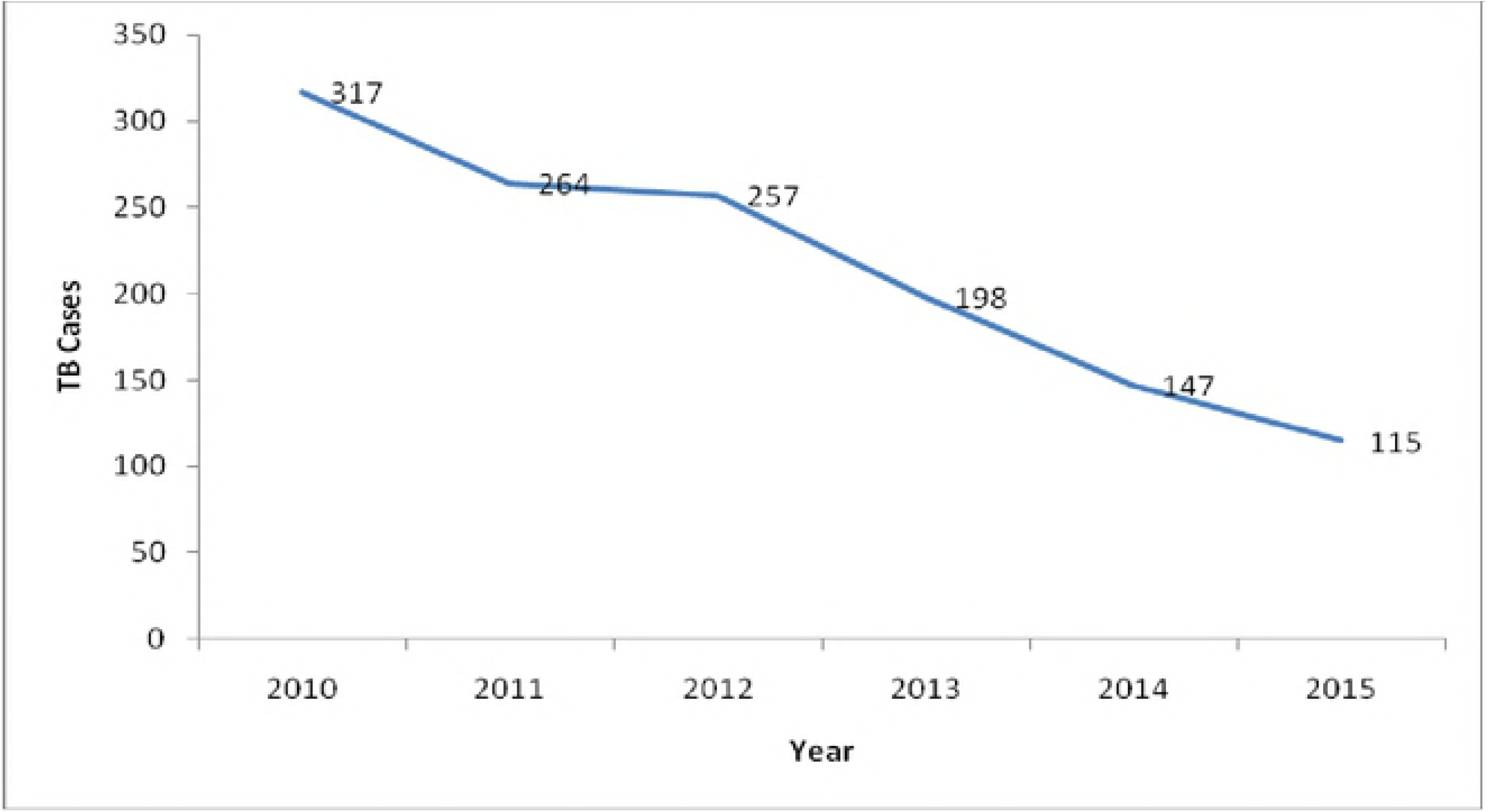
The number of tuberculosis patients notified between 2010-2015 by Zimbabwe Defence Forces health facilities.

**Fig 2.**
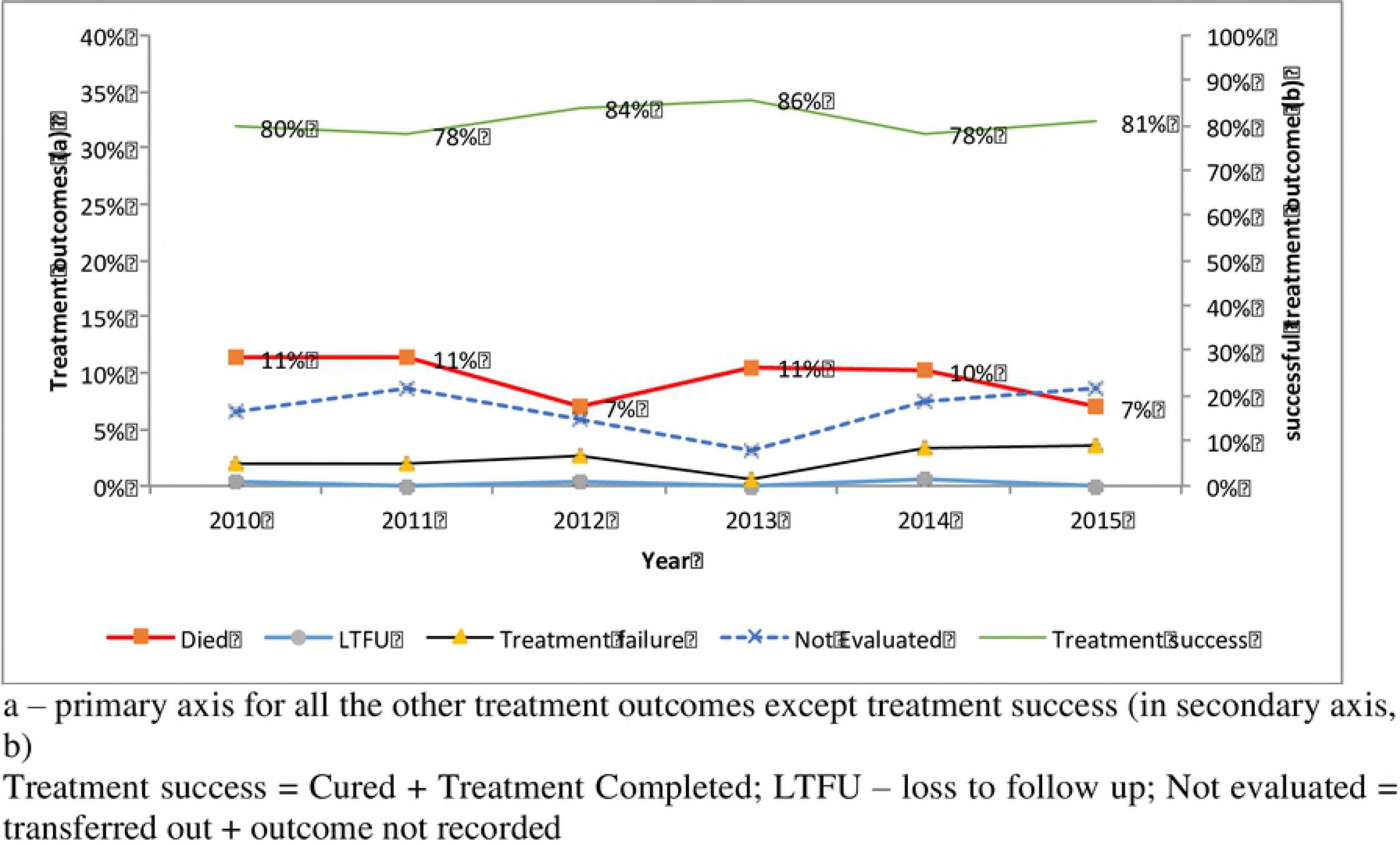
Trends in Tuberculosis treatment outcome between 2010-2015 in the Zimbabwe Defence Forces health facilities.

**Table 2.** Overall treatment outcomes among patients with tuberculosis managed in the Zimbabwe Defence Forces health facilities, 2010-2015.

| Treatment outcome | N | (%) |
| --- | --- | --- |
| Treatment success | 1053 | (81.1) |
| Treatment failure | 28 | (2.2) |
| Lost to follow-up | 3 | (0.2) |
| Died | 128 | (9.9) |
| Not evaluated | 86 | (6.6) |

HIV status and being on ART were significant factors associated with unfavourable TB treatment outcome, both before and after adjusting for covariates (**Table 3**). Compared to those HIV negative, those HIV co-infected and on ART were at 76% higher risk of experiencing unfavourable outcomes (95% CI: 1.25-2.45, *P*<0.001) while those not on ART were 3.5 times at higher risk (95% CI: 2.20-6.59, *P*<0.001). Sex, age, type and classification of TB were not associated with unfavourable treatment outcomes.

**Table 3.**
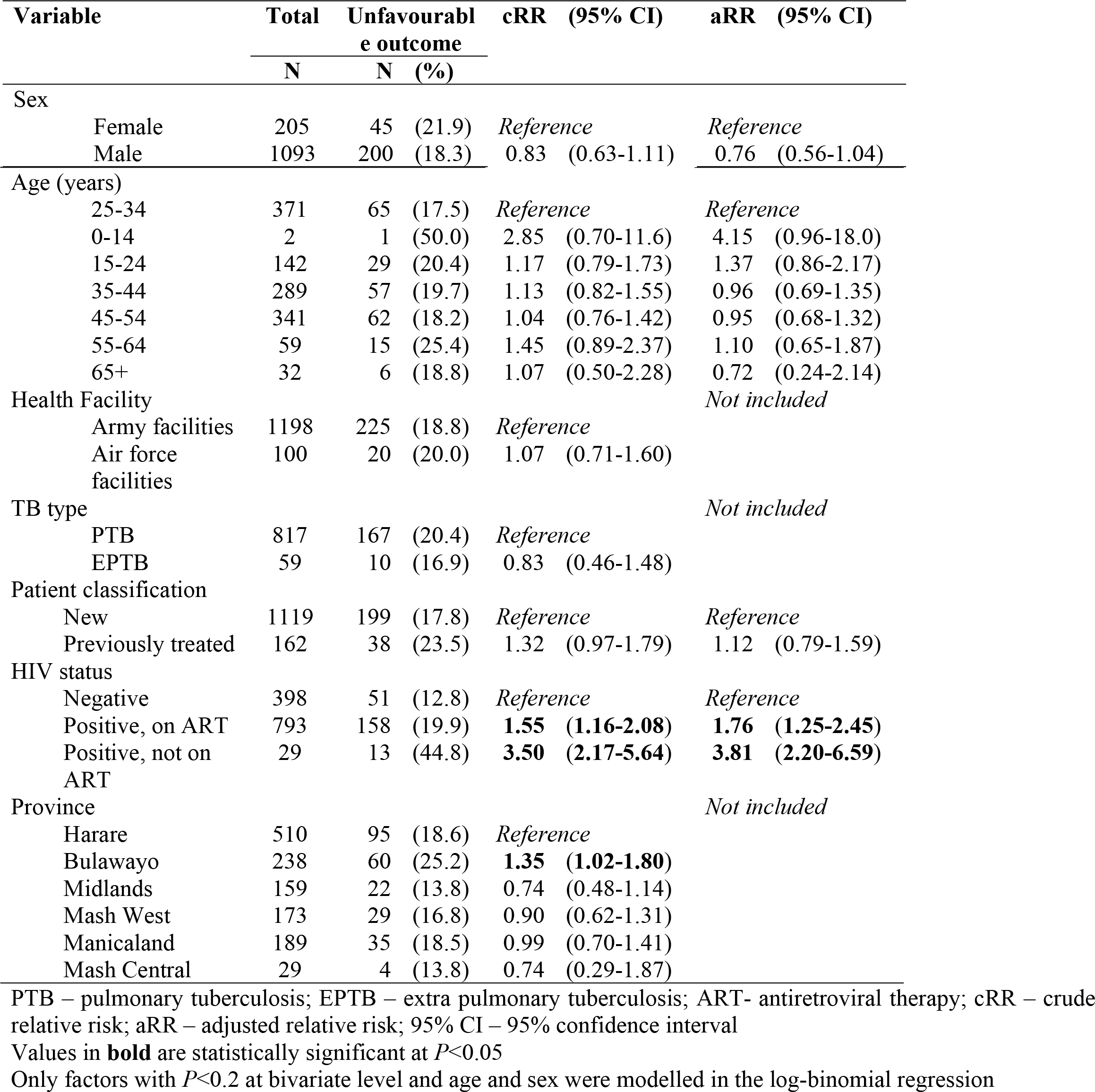
Factors associated with unfavourable tuberculosis treatment outcomes in the Zimbabwe Defence Forces health facilities, 2010-2015

## Discussion

This is the first study looking at TB notification and treatment outcomes in a defence force setting from Africa. The study established a two-fold reduction in TB notification over the six-year period while the treatment outcomes remained relatively stable. Treatment success rates were comparably high (81%) but mortality rates were also high, one-in-ten of notified cases, contributing to most of the unfavourable treatment outcomes. HIV co-infection and lack of ART were significantly associated with unfavourable TB treatment outcome. Those HIV co-infected on ART still had close to twice the risk of unfavourable outcome while those not on ART had four-times the risk, as compared to HIV negative TB patients.

The primary strength of our study was that we reviewed all TB patients notified in both Army and Airforce health facilities within Zimbabwe over six years. However, current TB clinical management documentation does not capture whether patients were military personnel or a dependent, which was a significant limitation. The impact of this lack of classification is likely low, because the majority of patients (>85%) were adult males. Additionally, 7% of patients did not have outcome data as they were not evaluated. It is suspected these patients actually transferred out, with care being provided elsewhere, but remains uncertain.

Another major challenge faced with interpreting our study results is the lack of prevalence data. To better assess the frequency of TB among the ZDF, particularly by geographical distribution, knowing the baseline military population would have been extremely valuable. This likely could have provided more meaningful insights to improving case notification and treatment outcomes. However, this was not possible because of security reasons and will likely always be difficult when studying this population.

Treatment success rates were comparable to the Zimbabwe averages, but are below the current WHO recommended goal of 90% success (2). These findings imply that the TB program within the ZDF – an at risk population – is at least on par with the country’s performance. This is admirable on several levels. First, military personnel are at increased risk for communicable diseases because of frequently living in confined, congested settings. Secondly, operational movements can make continuity of care challenging. Lastly, medication compliance could be difficult in this context and patient population.

The declining trend of TB in the Zimbabwe military is particularly noteworthy. MacPherson *et al* has previously published a declining trend of TB in Zimbabwe that was largely due to increased uptake of ART(5). This is likely the case with the ZDF, leading to the decreasing trend. HIV testing and ART uptake in this population was high at over 95%. This is remarkable and suggests that the quality of care for those PLWHIV within the military is excellent and without stigma. These findings should serve as a benchmark for other armed forces health services to strive for in the future.

Equally reassuring was that TB outcomes were similar, in both Army and Airforce health facilities. Likewise, outcomes were essentially the same across all facilities geographically, with the exception of Bulawayo, which has the highest HIV/TB prevalence in the country (ref). This suggests that the quality of TB care is equitable for armed services personnel and their dependents, whether they are based in the capital of Harare or elsewhere. Again, another benchmark the NTP program has set for others to follow.

The high death rates among the ZDF are notable. The 10% mortality rate may have been directly related to the extremely low LTFU (0.2%). Studies have previously shown that a considerable proportion of those classified as LTFU actually died(5). Similar to HIV care for those attending the ZDF facilities, TB quality of care and follow up were likely quite high. Nevertheless, some classified as not evaluated (including transfer outs) may actually have been LTFU.

Higher unfavourable treatment outcomes were still recorded even with ART, necessitating the need to establish the time to ART among the co-infected. In deed earlier ART has been shown to more beneficial in improving treatment outcomes compared to later ART (11) STRIDE, SAPIT studies. National TB programmes are encouraged to shift to monitoring and reporting of earlier ART (within 8 weeks of TB treatment) as opposed to ART within any time of TB treatment, and in line with WHO 2017 update (12).

Our study highlights both the successes and challenges the ZDF faces in TB case notification, treatment and follow up. We recommend that the NTP work with the ZDF health services to develop a more armed forces-specific monitoring tool going forward to improve outcome evaluation. Further research needs to be completed in the future focusing on identifying military personnel specifically without co-mingling dependents. Lastly, continued reporting on a regular basis by the ZDF will allow others to apply their lessons learned to improve care for military forces in similar contexts elsewhere.

## Conclusions

This study found decreasing trends of TB within the Zimbabwe Defence Forces from 2010-2015 with comparable treatment outcomes across the period. Although favourable TB treatment outcomes were high, they were still not at the current Zimbabwe National TB Programme and WHO goals. Continued monitoring of TB patients and evaluation of the TB programme is needed so as to attain higher treatment success rates and reduce deaths associated with TB.

## Acknowledgements

This research was conducted through the Structured Operational Research and Training Initiative (SORT IT), a global partnership led by the Special Programme for Research and Training in Tropical Diseases at the World Health Organization (WHO/TDR). The training model is based on a course developed jointly by the International Union Against Tuberculosis and Lung Disease (The Union) and Medécins sans Frontières (MSF). The specific SORT IT program which resulted in this publication was implemented by the Centre for Operational Research, The Union, Paris, France. Mentorship and the coordination/facilitation of this particular SORT IT workshop was provided through the Centre for Operational Research, The Union, Paris, France; the Department of Tuberculosis and HIV, The Union, Paris, France; the University of Washington, School of Public Health, Department of Global Health, Seattle, Washington, USA; and AMPATH, Eldoret, Kenya.

## Supporting Information

**S1** TB TREATMENT NOTIFICATIONS AND OUTCOMES

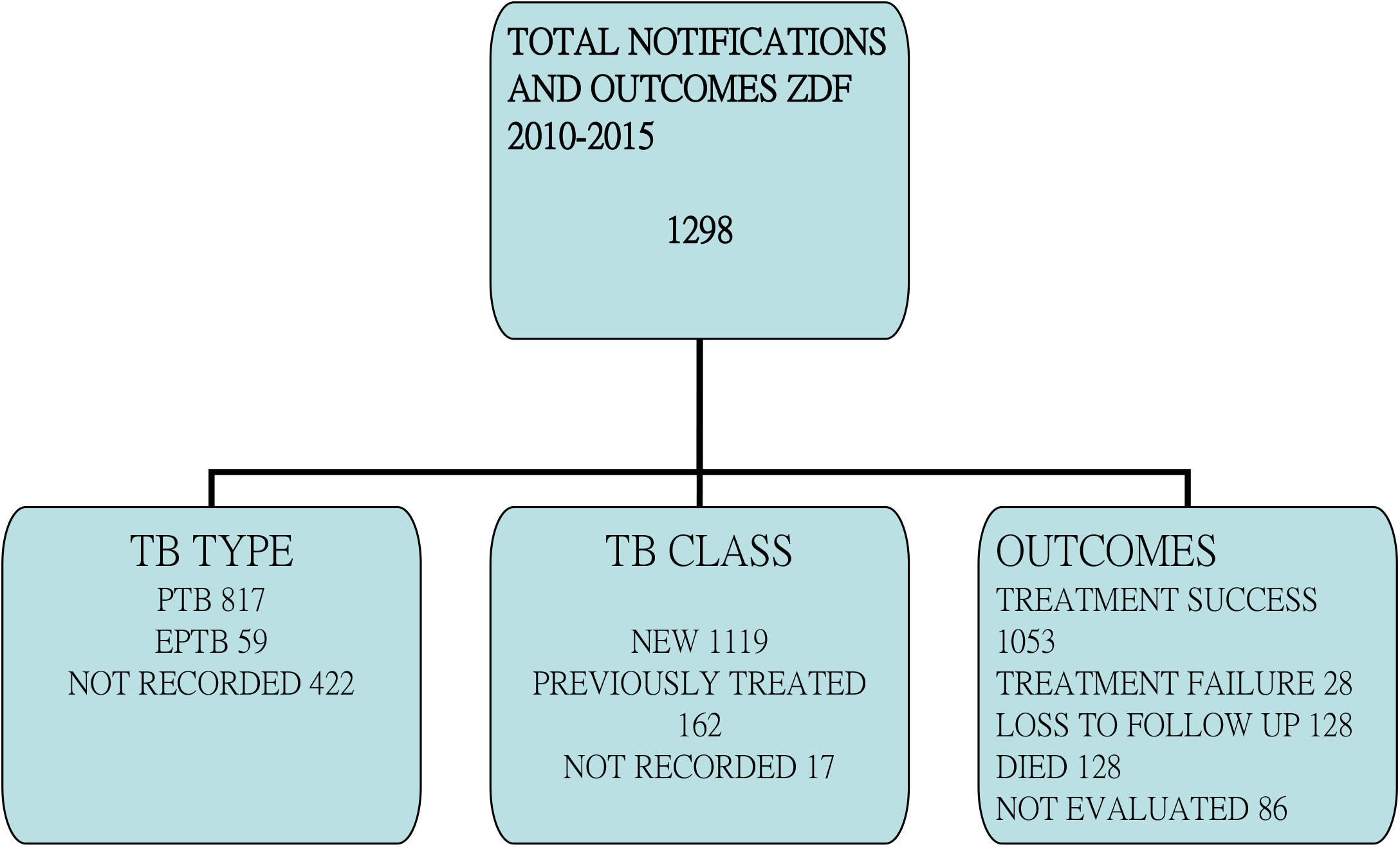

